# Chimeric antigen receptors that trigger phagocytosis

**DOI:** 10.1101/316323

**Authors:** Meghan A. Morrissey, Adam P. Williamson, Adriana M. Steinbach, Edward W. Roberts, Nadja Kern, Mark B. Headley, Ronald D. Vale

## Abstract

Chimeric antigen receptors (CARs) are synthetic receptors that reprogram T cells to kill cancer. The success of CAR-T cell therapies highlights the promise of programmed immunity, and suggests that applying CAR strategies to other immune cell lineages may be beneficial. Here, we engineered a family of Chimeric Antigen Receptors for Phagocytosis (CAR-Ps) that direct macrophages to engulf specific targets, including cancer cells. CAR-Ps consist of an extracellular antibody fragment, which can be modified to direct CAR-P activity towards specific antigens. By screening a panel of engulfment receptor intracellular domains, we found that the cytosolic domains from Megf10 and FcRγ robustly triggered engulfment independently of their native extracellular domain. We show that CAR-Ps drive specific engulfment of antigen-coated synthetic particles and whole cancer cells. Addition of a tandem PI3K recruitment domain increased cancer cell engulfment. Finally, we show that CAR-P expressing macrophages reduce cancer cell number in co-culture by over 40%.

**Summary:** We report the first Chimeric Antigen Receptors for Phagocytosis (CAR-Ps) that promote engulfment of antigen-coated particles and cancer cells.

## Introduction

Chimeric antigen receptors (CARs) are synthetic transmembrane receptors that redirect T cell activity towards clinically relevant targets (reviewed in (Lim et al. 2017; Fesnak, June, and Levine 2016)). The CAR-T receptor contains an extracellular single chain antibody fragment (scFv) that recognizes known tumor antigens, and intracellular signaling domains from the T Cell Receptor (TCR) and costimulatory molecules that trigger T cell activation (Fesnak, June, and Levine 2016; Kochenderfer et al. 2009). CAR-T cells recognizing CD19, a marker expressed at high levels on the surface of B cells and B cell-derived malignancies, have been used successfully to target hematological malignancies with 70-90% of patients showing measurable improvement (Lim et al. 2017; Engel et al. 1995; Haso et al. 2013). The success of CAR-T suggests that programming immune cells to target cancer might be a broadly applicable approach.

Macrophages are critical effectors of the innate immune system, responsible for engulfing debris and pathogens. Harnessing macrophages to combat tumor growth is of longstanding interest (C. Alvey and Discher 2017; Lee et al. 2016). Macrophages are uniquely capable of penetrating solid tumors, while other immune cells, like T cells, are physically excluded or inactivated (Lim et al. 2017; Lee et al. 2016). This suggests that engineered macrophages may augment existing T cell-based therapies. Early efforts transferring healthy macrophages into cancer patients failed to inhibit tumor growth, suggesting that macrophages require additional signals to direct their activity towards tumors (Lacerna, Stevenson, and Stevenson 1988; Andreesen et al. 1990). Antibody blockade of CD47, a negative regulator of phagocytosis, reduced tumor burden, indicating that shifting the balance in favor of macrophage activation and engulfment is a promising therapeutic avenue (Majeti et al. 2009; Chao et al. 2010; Jaiswal et al. 2009; Tseng et al. 2013). Here, we report a family of chimeric antigen receptors that activate phagocytosis of cancer cells based on recognition of defined cell surface markers, resulting in significantly reduced cancer cell growth.

## Results

To program engulfment towards a target antigen, we created a CAR strategy using the CAR-T design as a guide (Fesnak, June, and Levine 2016). We call this new class of synthetic receptors Chimeric Antigen Receptors for Phagocytosis (CAR-Ps). The CAR-P molecules contain the extracellular single-chain antibody variable fragment (scFv) recognizing the B cell antigen CD19 (αCD19) and the CD8 transmembrane domain present in the αCD19 CAR-T (Fesnak, June, and Levine 2016; Kochenderfer et al. 2009). To identify cytoplasmic domains capable of promoting phagocytosis, we screened a library of known murine phagocytic receptors: Megf10 (Fig. 1a), the common γ subunit of Fc receptors (FcRγ), Bai1, and MerTK (Penberthy and Ravichandran 2016). FcR triggers engulfment of antibody-bound particles, while the other receptors recognize apoptotic corpses (Freeman and Grinstein 2014; Penberthy and Ravichandran 2016). We also made a receptor containing an extracellular αCD19 antibody fragment and a cytoplasmic GFP, but no signaling domain, to test whether adhesion mediated by the αCD19 antibody fragment is sufficient to induce engulfment (Fig. 1a; CAR-P^GFP^).

**Figure 1:**
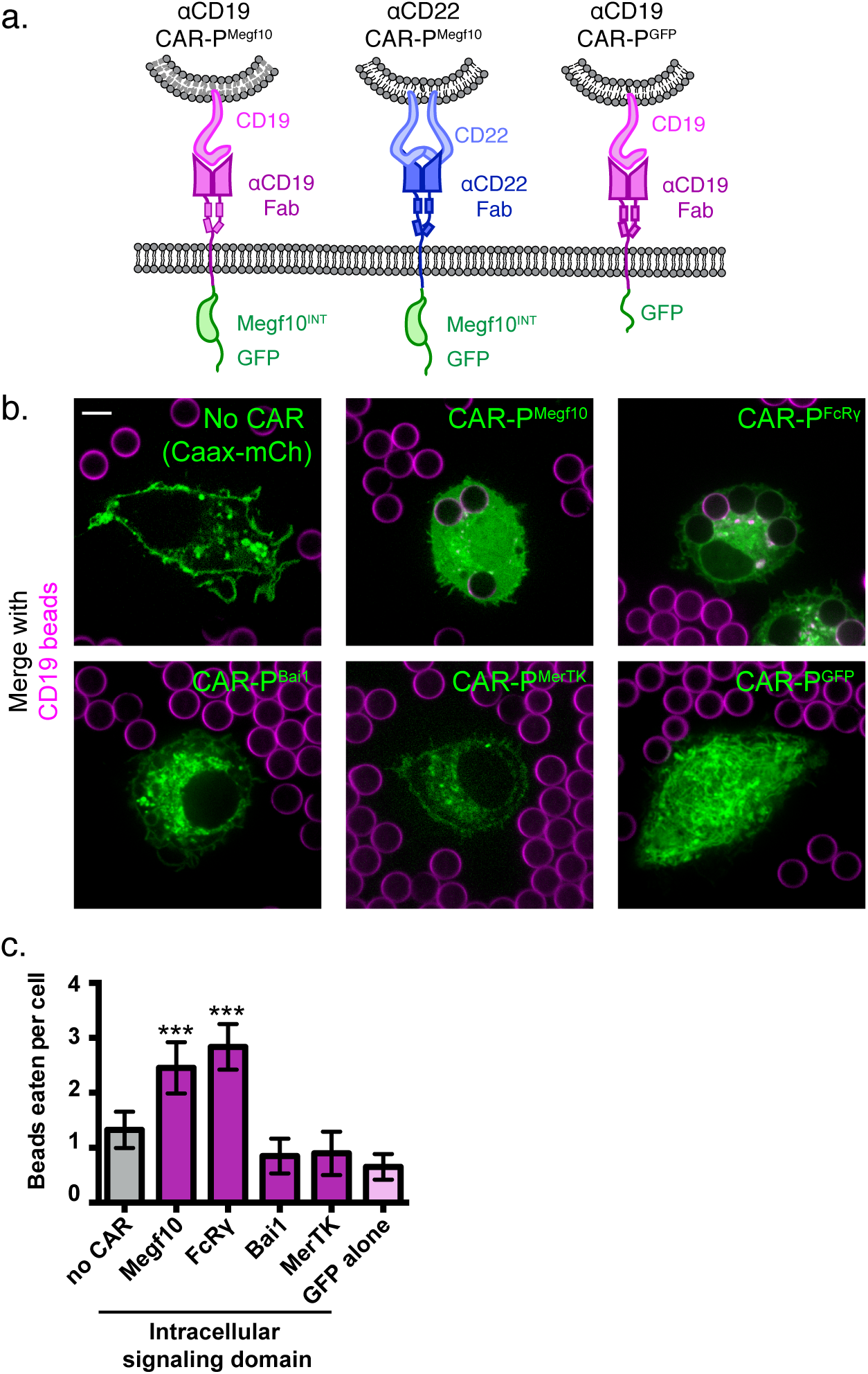
Identification of intracellular signaling region for CAR-P. (A) Schematics show the structure of CAR-P constructs. An αCD19 (purple) or αCD22 (blue, center) scFv directs CAR specificity. Intracellular signaling domains from Megf10 or the indicated engulfment receptor (green) activate engulfment. CAR-P^GFP^ contains only GFP and no intracellular signaling domains (right). All constructs include a transmembrane domain from CD8 and a C-terminal GFP. (B) J774A.1 macrophages expressing αCD19 CAR-P with the indicated intracellular signaling domain (green) engulf 5 μm silica beads covered with a supported lipid bilayer containing His-tagged CD19 extracellular domain. The beads were visualized with atto390-labeled lipid incorporated into the supported lipid bilayer (magenta). Cells infected with the cell membrane marker, mCherry-CAAX, were used as a control (no CAR, top left). The average number of beads eaten per cell is quantified in (C). The scale bar indicates 5 μm and n=78–163 cells per condition, collected during three separate experiments. Error bars denote 95% confidence intervals and *** indicates p<0.0001 compared to mCherry-CAAX control by Kruskal-Wallis test with Dunn’s multiple comparison correction.

To assay our library of CAR-Ps, we introduced each CAR-P into J774A.1 macrophages by lentiviral infection. As an engulfment target, we used 5 μm diameter silica beads coated with a supported lipid bilayer. A His_8_-tagged extracellular domain of CD19 was bound to a NiNTA-lipid incorporated into the supported lipid bilayers. Macrophages expressing a CAR-P with the Megf10 (CAR-P^Megf10^) or FcRγ (CAR-P^FcRγ^) intracellular domain promoted significant engulfment of CD19 beads compared to macrophages with no CAR (Fig. 1b, c; Video 1). Macrophages expressing CAR-P^Bai1^, CAR-P^MerTK^, and the adhesion-only CAR-P^GFP^ did not bind the CD19 beads even though these CAR-Ps are present at the cell surface (Fig. 1b, c; Supplementary Fig. 1).

Next we asked if the CAR-P strategy could target a different antigen. Because CAR- P^Megf10^ performed well in our initial screen, we developed αCD22 CAR-P^Megf10^ using a previously developed αCD22 antibody fragment (Fig. 1a) (Xiao et al. 2009; Haso et al. 2013). Consistent with our results using αCD19-based CARs, αCD22 CAR-P^Megf10^ promoted engulfment of CD22 beads (Fig. 2a). To confirm antigen specificity of CAR-P, we incubated αCD19 CAR-P^Megf10^ macrophages with CD22 beads, and αCD22 CAR- P^Megf10^ macrophages with CD19 beads. CD19 beads were not eaten by αCD22 CAR- P^Megf10^ macrophages, and CD22 beads were not eaten by αCD19 CAR-P^Megf10^ macrophages (Fig. 2a). These data indicate that CAR-P^Megf10^ specifically triggers engulfment in response to the target ligand and that the CAR-P strategy is able to target multiple cancer antigens.

**Figure 2:**
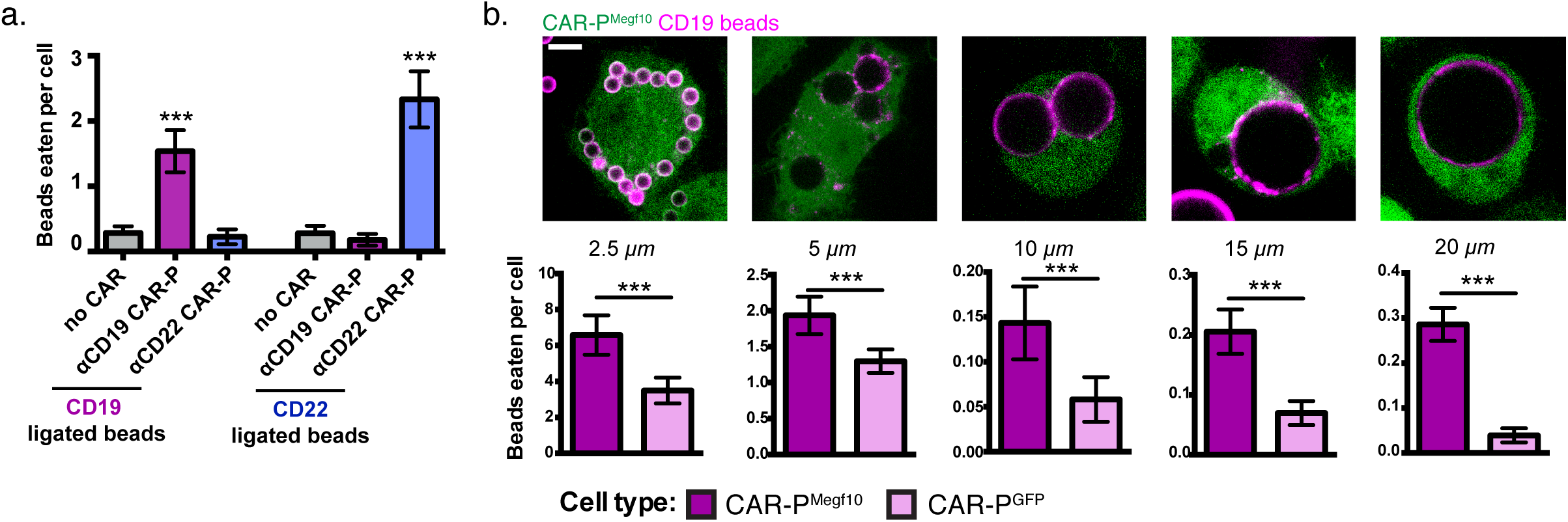
CAR-P expression drives specific engulfment of diverse beads. (A) Macrophages infected with the αCD19 (purple) or αCD22 (blue) CAR-P^Megf10^ or mCherry-CAAX control were fed either CD19-ligated beads (left) or CD22-ligated beads (right). Engulfment is quantified as the mean beads eaten per cell (n=120–157 cells per condition collected during three separate experiments). Error bars denote 95% confidence intervals and *** indicates p<0.0001 compared to mCherry-CAAX control by Kruskal-Wallis test with Dunn’s multiple comparison correction. (B) J774A.1 macrophages expressing the αCD19 CAR-P^Megf10^ (green) were fed beads of various sizes (magenta, diameter of bead indicated below image). All beads are covered in a supported lipid bilayer, with His-tagged CD19 extracellular domain ligated to the bilayer. The frequency of engulfment is quantified below (n=169–760 cells per condition, pooled from at least three separate experiments). Error bars denote 95% confidence intervals of the mean. *** indicates p<0.0001 respectively by Mann-Whitney test. All scale bars represent 5 *μ*m.

To further define the capabilities of the CAR-P, we assessed the capacity of CAR-P-expressing macrophages to engulf variably sized targets. We found that CAR-P^Megf10^ was able to trigger specific engulfment of beads ranging from 2.5 μm to 20 μm in diameter, with higher specificity above background engulfment being demonstrated for the larger beads (Fig. 2b).

To determine if the CAR-P^Megf10^ initiates active signaling at the synapse between the macrophage and target, we stained for phosphotyrosine. Macrophages expressing CAR-P^Megf10^ exhibited an increase in phosphotyrosine at the synapse, while macrophages expressing CAR-P^GFP^ did not show this enrichment (Fig. 3a). This result suggests that CAR-P^Megf10^ initiates engulfment through a localized signaling cascade involving tyrosine phosphorylation.

**Figure 3:**
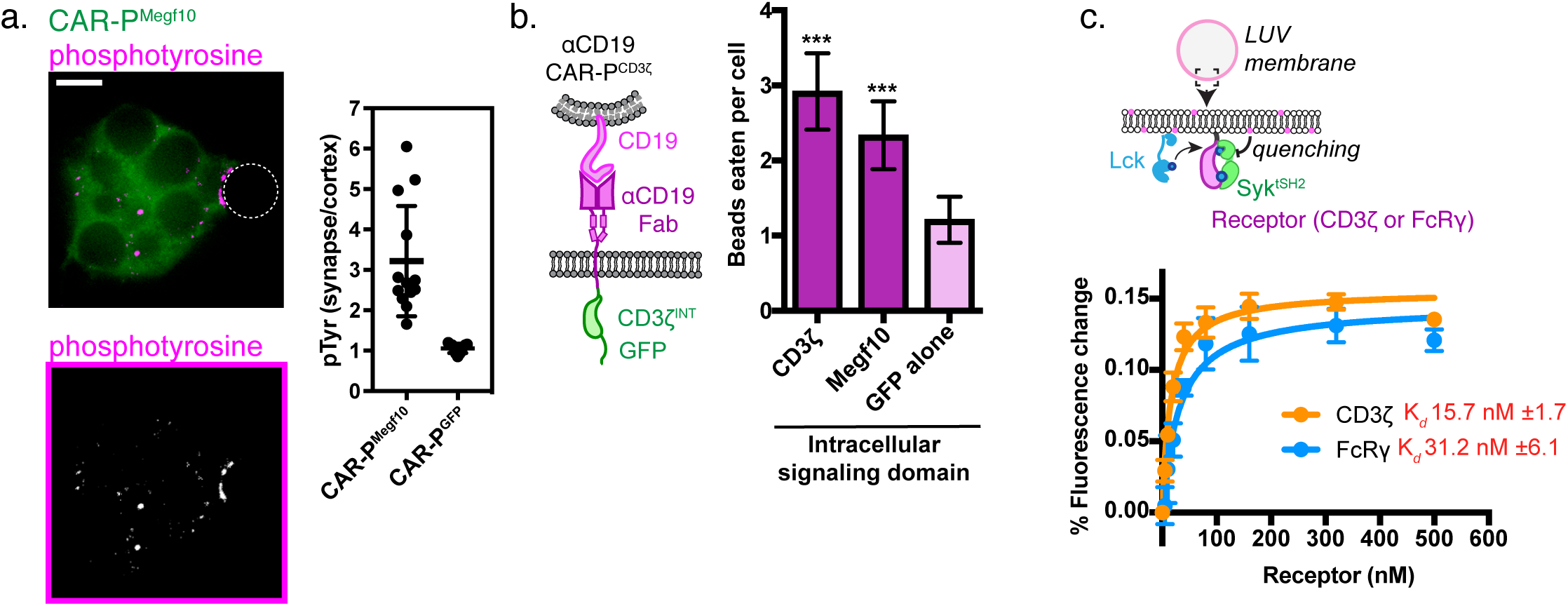
A phosphorylated ITAM at the cell-target synapse drives engulfment. (A) Macrophages expressing αCD19 CAR-P^Megf10^ (green, top) or αCD19 CAR-P^GFP^ were incubated with CD19-ligated beads (position indicated with dotted line), fixed and stained for phosphotyrosine (magenta, top; greyscale, bottom). The fold enrichment of phosphotyrosine at the cell-bead synapse compared to the cell cortex is graphed on the right (n≥11; each dot represents one cell-bead synapse; lines represent the mean ± one standard deviation). (B) Schematic shows the structure of CAR-P constructs in the plot at right. An αCD19 (purple) scFv directs CAR specificity. The intracellular signaling domains from CD3ζ activates engulfment. At right, J774A.1 macrophages expressing the αCD19 with the indicated intracellular signaling domain engulf 5 μm silica beads covered with a supported lipid bilayer containing His-tagged CD19 extracellular domain. The average number of beads eaten per cell is quantified (n=156–167 per condition collected during three separate experiments). Error bars denote 95% confidence intervals and *** indicates p<0.0001 compared to CAR-P^GFP^ control by Kruskal-Wallis test with Dunn’s multiple comparison in correction. (C) Model of the liposome-based fluorescence quenching assay used to determine affinity between the Syk tSH2 domains and the receptor tails of CD3ζ and FcRγ, two intracellular signaling domains that promote engulfment. Binding between the Syk tSH2 reporter (Syk tSH2), green, and a receptor tail, purple, was detected by rhodamine quenching of BG505 dye on the reporter (see Methods). Kd was determined by assessing mean fluorescence quenching for the last 20 timepoints collected ~45 min after ATP addition over a receptor titration from 0 to 500 nM. Each point represents the mean ± SD from 3 independent experiments. Kd ± SE was calculated by nonlinear fit assuming one site specific binding.

Both successful CAR-P intracellular domains (from FcRγ and Megf10) have cytosolic Immunoreceptor Tyrosine-based Activation Motifs (ITAMs) that are phosphorylated by Src family kinases. Based on this observation, we hypothesized that the expression of an alternate ITAM-containing receptor might initiate phagocytosis when expressed in macrophages. The CD3ζ subunit of the T cell receptor contains three ITAM motifs. To test if the CD3ζ chain was able to activate phagocytic signaling, we transduced macrophages with the first generation CAR-T (Fig. 3b). The CAR-T was able to trigger engulfment of CD19 beads to a comparable extent as CAR-P^Megf10^ (Fig. 3b). In T cells, phosphorylated ITAMs in CD3ζ bind to tandem SH2 domains (tSH2) in the kinase ZAP70. Zap70 is not expressed in macrophages, but Syk, a phagocytic signaling effector and tSH2 domain containing protein, is expressed at high levels (Andreu et al. 2017). Previous work suggested that Syk kinase can also bind to the CD3ζ ITAMs (Bu, Shaw, and Chan 1995), indicating that the CAR-T may promote engulfment through a similar mechanism as CAR-P^FcRγ^. To quantitatively compare the interaction between Syk^tSH2^ and CD3ζ or FcRγ in a membrane proximal system recapitulating physiological geometry, we a used liposome-based assay (Fig. 3c (Hui and Vale 2014)). In this system, His_10_-CD3ζ and His_10_-Lck (the kinase that phosphorylates CD3ζ) are bound to a liposome via NiNTA-lipids and the binding of labeled tandem SH2 domains to phosphorylated CD3ζ was measured using fluorescence quenching. Our results show that Syk-tSH2 binds the CD3ζ and FcRγ with comparable affinity (~15 nM and ~30 nM respectively, Fig. 3c). Collectively, these results demonstrate that the TCR CD3ζ chain can promote phagocytosis in a CAR-P, likely through the recruitment of Syk kinase.

We next sought to program engulfment towards a cellular target. We incubated the CAR-P^Megf10^ and CAR-P^FcRγ^ macrophages with cancerous Raji B cells that express high levels of endogenous CD19. Strikingly, the majority of CAR-P-expressing macrophages internalized bites of the target cell (Fig. 4a, Video 2, 78% of CAR-P^Megf10^ and 85% of CAR-P^FcRγ^ macrophages internalized bites within 90 min). The biting phenotype resembles trogocytosis, or nibbling of live cells, which has been reported previously in immune cells (Joly and Hudrisier 2003). This process was dependent on the ITAM-bearing intracellular signaling domain, as removing the signaling domain (CAR-P^GFP^) dramatically reduced trogocytosis (Fig. 4a). Enrichment of phosphotyrosine at the cell-cell synapse further supports active signaling initiating trogocytosis (Supplementary Fig. 2). The CAR-P module also was able to induce trogocytosis in non-professional phagocytes, NIH 3T3 fibroblast cells (Supplementary Fig. 3). This suggests that the CAR-P can promote cancer antigen-dependent engulfment by both professional and non-professional phagocytes.

**Figure 4:**
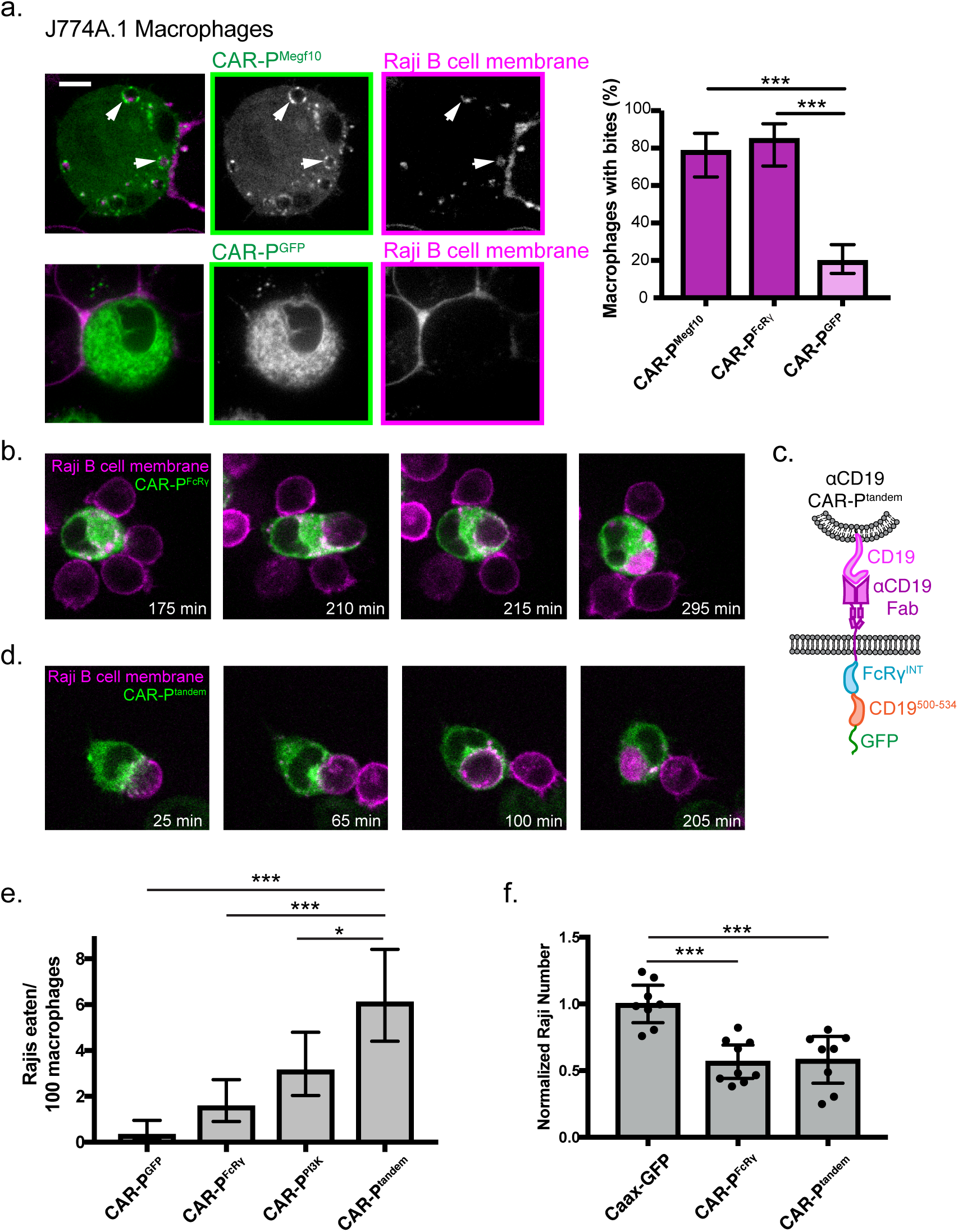
CAR-P promotes trogocytosis and whole cell eating. (A) J774A.1 macrophages expressing the αCD19 CAR-P^Megf10^ (top panel, green in merge, left; greyscale, center) engulf pieces of CD19+ Raji B cells (labeled with mCherry-CAAX; magenta in merge, left; greyscale, right). The corresponding control αCD19 CAR-P^GFP^-infected cells are shown below. Arrows point to pieces of ingested Raji B cell. The proportion of CAR-P expressing macrophages internalizing one or more bite within 90 min is quantified on the right. Bites are defined as a fully inter-nalized mCherry-positive vesicle >1 μm in diameter; n=46 CAR-P^Megf10^ macrophages, n=39 CAR-P^FcRγ^ macrophages and 102 CAR-P^GFP^ macrophages acquired during three separate experiments. (B) Time course of a J774A.1 macrophage expressing CAR-P^FcRγ^ (green) internalizing a whole Raji B cell labeled with mCherry-CAAX (magenta). These images correspond to frames from Video 3. (C) Schematic shows the structure of CAR-P^tandem^ construct, combining the intracellular signaling domain from FcRγ and the p85 recruitment domain from CD19. (D) Time course of a J774A.1 macrophage expressing CAR-P^tandem^ (green) internalizing a whole Raji B cell labeled with mCherry-CAAX (magenta). These images correspond to frames from Video 4. (E) Macrophages and Raji B cells were incubated together at a 1:2 macrophage:Raji ratio, and the number of whole Raji B cells eaten per 100 macrophages during 4–8 hr of imaging is graphed. Graph depicts pooled data from 4 independent experiments; n = 921 CAR-P^GFP^, n = 762 CAR-P^FcRγ^, n = 638 CAR-P^Pi3K^, n = 555 CAR-P^tandem^. Sample sizes were selected for their ability to detect a 5% difference between samples with 95% confidence. (F) 10,000 macrophages and 20,000 Raji B cells were incubated together for 44 hr. The number of Rajis was then quantified by FACS. 2–3 technical replicates were acquired each day on three separate days. The number of Rajis in each replicate was normalized to the average number present in the CAAX-GFP macrophage wells on that day. * indicates p<0.01, *** indicates p<0.0001 by two-tailed Fisher Exact Test (a and e) or by Ordinary one way ANOVA with Dunnet’s correction for multiple comparisons (f); error bars denote 95% confidence intervals.

We next focused on engineering strategies to engulf whole cancer cells. We observed that macrophages expressing the CAR-P^Megf10^ or CAR-P^FcRγ^ were capable of engulfing whole cells (2 cancer cells eaten per 100 macrophages in a 4-8 hr window for both CAR-P^Megf10^ or CAR-P^FcRγ^, Fig 4b,e, Video 3). Whole cell engulfment was infrequent but trogocytosis was robust, suggesting that even productive macrophage target interactions were frequently insufficient to trigger whole cell engulfment. We hypothesized that combining signaling motifs in a tandem array might increase the frequency of whole cell engulfment by specifically recruiting effectors required for the engulfment of large targets. Previous work demonstrated that PI3K signaling is important for internalization of large targets (Schlam et al. 2015). To increase PI3K recruitment to the CAR-P, we fused the portion of the CD19 cytoplasmic domain (amino acids 500 to 534) that recruits the p85 subunit of PI3K to the CAR-P^FcRγ^ creating a "tandem” CAR (CAR-P^tandem^, Fig 4c) (Tuveson et al 1993, Brooks et al 2004). A CAR-P containing the p85 recruitment motif alone (CAR-P^PI3K^) was able to induce some whole cell engulfment, comparable to the CAR-P^FcRγ^ (Fig 4e). Expression of CAR-P^tandem^ tripled the ability of macrophages to ingest whole cells compared to CAR-P^GFP^ (6 cancer cells eaten per 100 macrophages, Fig. 4d,e, Video 4). These data indicate that assembling an array of motifs designed to recruit distinct phagocytic effectors can increase CAR-P activity towards whole cells.

To determine if the combination of whole cell eating and trogocytosis was sufficient to drive a noticeable reduction in cancer cell number, we incubated CAR-P macrophages with Raji B cells for two days. After 44 hr of co-culture, we found that CAR-P macrophages significantly reduced the number of Raji cells (Fig 4f). Although the CAR-P^tandem^ was much more efficient at whole cell eating, the CAR-P^FcRγ^ performed nearly as well at eliminating Rajis. Importantly, our assay does not distinguish between whole cell engulfment or death following trogocytosis, so it is possible both CAR-P activities are contributing to Raji death rates. Overall, this data suggests that the CAR-P is a successful strategy for directing macrophages towards cancer targets, and can initiate whole cell eating and trogocytosis leading to cancer cell elimination.

In summary, we engineered phagocytes that recognize and ingest targets through specific antibody-mediated interactions. This strategy can be directed towards multiple extracellular ligands (CD19 and CD22) and can be used with several intracellular signaling domains that contain ITAM motifs (Megf10, FcRγ, and CD3ζ. Previous work has suggested that spatial segregation between Src-family kinases and an inhibitory phosphatase, driven by receptor ligation, is sufficient to trigger signaling by the T cell receptor (Davis and van der Merwe 2006; James and Vale 2012) and FcR (Freeman et al. 2016). The CAR-Ps that we have developed may similarly convert receptor-ligand binding into receptor phosphorylation of ITAM domains through partitioning of kinases and phosphatases at the membrane-membrane interface.

Further development of CAR-Ps could be useful on several therapeutic fronts. Targeting of tumor cells by macrophages has been suggested to cause tumor cell killing (Jaiswal et al. 2009; Majeti et al. 2009; Chao et al. 2010; Jadus et al. 1996), either through directly engulfing cancer cells or by stimulating antigen presentation and a T cell-mediated response (Liu et al. 2015; Tseng et al. 2013). Inhibition of the CD47-SIRPA "Don’t eat me” signaling pathway has also been shown to result in engulfment of cancer cells (Chen et al. 2017; Gardai et al. 2005; Jaiswal et al. 2009; Majeti et al. 2009; Chao et al. 2010). A recent study suggests that CD47 inhibition is most effective when combined with a positive signal to promote target engulfment, which raises the possibility of combining CAR-P expression with CD47 or SIRPA inhibition for an additive effect (Alvey et al. 2017).

Although we were able to increase whole cell engulfment by recruiting the activating subunit of PI3K to the phagocytic synapse, the engulfment of larger 20 micron beads was more frequent than the engulfment of whole cells. We hypothesize that this is due to differing physical properties of the engulfment target. Specifically, increased target stiffness has been shown to promote engulfment, suggesting that manipulating the physical properties of the engulfment target could also be a potential strategy for increasing CAR-P efficiency (Beningo and Wang 2002; Cross et al. 2007).

While the CAR-P can engulf whole, viable cancer cells, the ingestion of a piece of the target cell is more common. Trogocytosis, or nibbling of living cells, has also been described in immune cells (Harshyne et al. 2003; Harshyne et al. 2001; Kao et al. 2006; Joly and Hudrisier 2003; Batista, Iber, and Neuberger 2001) and brain-eating amoebae (Ralston et al. 2014). In vivo studies also have shown that endogenous dendritic cell populations ingest bites of live tumor cells, contributing to presentation of cancer neo-antigen (Harshyne et al. 2001; Harshyne et al. 2003). Although we were able to use the CAR-P to induce trogocytosis in dendritic cells, we were not able to detect robust cross presentation of the model antigen ovalbumin (Fig S4). Thus, although using CAR-Ps to enhance cross presentation of cancer antigen is an intriguing future avenue, such a strategy would likely require more optimization of the dendritic cell subset employed or the CAR-P receptor itself.

Overall, our study demonstrates that the CAR approach is transferrable to biological processes beyond T cell activation and that the expression of an engineered receptor in phagocytic cells is sufficient to promote specific engulfment and elimination of cancer cells.

## Acknowledgments

We thank J. Reiter and E. Yu for providing mouse long bones as a source for hematopoetic stem cells. We thank K. McKinley and X. Su for critical feedback on this manuscript. M.A.M. was supported by the National Institute of General Medical Sciences of the National Institutes of Health under award number F32GM120990. A.P.W was supported by a CRI Irvington Postdoctoral Fellowship. This work was funded by the Howard Hughes Medical Institute.

## Materials and Methods

### Constructs and Antibodies

Detailed information for all constructs can be found in Supplemental Excel File 1, “Constructs” tab. This file includes the following information for all receptors developed in this study: signal peptide, extracellular antibody fragment, stalk/transmembrane domain, and cytosolic tail including appropriate accession numbers. Antibodies used in this study are described in Supplemental Excel File 1, “Antibodies” tab.

### Cell culture

J774A.1 macrophages and NIH 3T3 fibroblasts were obtained from the UCSF cell culture facility and cultured in DMEM (Gibco, Catalog #11965–092) supplemented with 1 x Pen-Strep-Glutamine (Corning, Catalog #30–009-Cl) and 10% fetal bovine serum (FBS) (Atlanta Biologicals, Catalog #S11150H). Raji B cells were obtained from J. Blau (McManus lab, UCSF) and cultured in RPMI (Gibco, Catalog #11875–093) supplemented with 1 x Pen-Strep-Glutamine (Corning, Catalog #30–009-Cl), 10% FBS (Atlanta Biologicals, Catalog #S11150H), 10mM HEPES (Gibco, Catalog #1530080), and 5 μM 2-Mercaptoethanol (Sigma, Catalog #M6250–100mL). All cell lines used in this study were tested for Mycoplasma at least once per month using the Lonza MycoAlert Detection Kit (Lonza, Catalog# LT07–318) and control set (Lonza, Catalog #LT07–518).

### Lentivirus production and infection

Lentiviral infection was used to stably express CAR-P constructs in all cell types. Lentivirus was produced by HEK293T cells transfected with pMD2.G (a gift from Didier Tronon, Addgene plasmid # 12259 containing the VSV-G envelope protein), pCMV-dR8.91 (since replaced by 2^nd^ generation compatible pCMV-dR8.2, Addgene plasmid #8455), and a lentiviral backbone vector containing the construct of interest (derived from pHRSIN-CSGW) using lipofectamine LTX (Invitrogen, Catalog # 15338–100). The media on the HEK293T cells was replaced with fresh media 8–16 hours post transfection to remove transfection reagent. At 50–72 hr post-transfection, the lentiviral media was filtered with a 0.45 μm filter and concentrated by centrifugation at 8,000 x g for 4 hr or overnight. The concentrated supernatant was applied directly to ~0.5 x 10^6^ NIH 3T3 cells in 2 ml of fresh media. For J77A4.1 macrophages and Raji B cells, the concentrated supernatant was mixed with 2 mls of media and 2 pg lipofectamine (Invitrogen, Catalog # 18324–012) and added to the cells. The cells were spun at 2,200 x g for 45 min at 37°C. Cells were analyzed a minimum of 72 hr later.

### Preparation of CD19 and CD22 5 μm silica beads

Chloroform-suspended lipids were mixed in the following molar ratio using clean glasstight Hamilton syringes (Hamilton, Catalog #8 1100): 97% POPC (Avanti, Catalog # 850457), 2% Ni2+-DGS-NTA (Avanti, Catalog # 790404), 0.5% PEG5000-PE (Avanti, Catalog # 880230, and 0.5% atto390-DOPE (ATTO-TEC GmbH, Catalog # AD 390–161). Lipid mixes were dried under argon and desiccated overnight under foil. Dried lipids were resuspended in 1 ml tissue-culture grade PBS, pH7.2 (Gibco, Catalog # 20012050), and stored under argon gas. Small unilamellar vesicles were formed by 5 freeze-thaw cycles followed by 2 × 5 min of bath sonication (Bioruptor Pico, Diagenode), and cleared by ultracentrifugation (TLA120.1 rotor, 35,000 rpm / 53,227 × g, 35 minutes, 4°C) or by 33 freeze thaw cycles. Lipid mixes were used immediately for form bilayers or shock frozen in liquid nitrogen and stored under argon at −80°C. To form bilayers on silica beads, 6×10^8^ 5 μm silica microspheres (10% solids, Bangs Labs, Catalog # SS05N) were washed 2x in water, and 2x in PBS by sequential suspension in water and spinning at 800 rcf, followed by decanting. Cleaned beads were resuspended in 150 μl tissue-culture grade PBS, pH7.2 (Gibco, Catalog # 20012050) and briefly vortexed. 30 μl cleared SUVs prepared as above as a 10 mM stock were added to bead suspension for a 2 mM final SUV concentration. Beads were vortexed for 10 sec, covered in foil, and rotated for 30 min at room temperature to form bilayers. Bilayer-coated beads were washed 3x in PBS by sequential centrifugation at 800 rcf and decanting. Beads were resuspended in PBS + 0.1% w/v BSA for blocking for 15 min rotating at room temperature under foil. 10 nM final concentration of CD19_his8_ (Sino Biological, Catalog # 11880H08H50) or CD22_his8_ (Sino Biological, Catalog # 11958H08H50) protein were added to blocked beads and proteins were allowed to bind during a 45 min incubation rotating under foil at room temperature. Beads were washed 3x in PBS + 0.1% w/v BSA by sequential centrifugation at 300 rcf and decanting. Beads were resuspended in 120 ml PBS + 0.1% w/v BSA.

### Preparation of CD19 silica beads over a range of diameters

Prior building bilayers on Silica beads ranging from 2.5 μm-20 μm in diameter (Microspheres-Nanospheres, Catalog# C-SIO-2.5, 5, 10, 15, 20), beads were RCA cleaned as follows: beads were pelleted at 2,000xg in low retention tubes (Eppendorf, Catalog #022431081) and resuspended in acetone. Resuspended beads were sonicated for 60 min in a bath sonicator. Rinse and sonication were repeated in ethanol. Finally, rinse and sonication were repeated in water. Beads were then washed 2x in water to remove all traces of ethanol and left in a small volume after decanting. All further steps were performed in a 70–80ΰC water bath prepared in a fume hood. Proper Personal Protective Equipment (PPE) was worn throughout the RCA cleaning protocol. Washed beads were added to 3 ml of hot 1.5 M KOH in a clean glass vial suspended in the water bath described above. 1 ml 30% H2O2 to bead solution and allowed to react for 10 minutes. Washed beads were cooled on ice, pelleted at 2,000xg and rinsed 5x in ultrapure water. Used cleaning solution was saved for disposal by Environmental Health & Safety (EH&S). Cleaned beads were resuspended in 240 μl tissue-culture grade PBS, pH7.2 (Gibco, Catalog # 20012050) and briefly vortexed. The lipid mix used in this assay differed slightly from above. Here a mix of 93.5% POPC (Avanti, Catalog # 850457), 5% Ni^2+-^DGS-NTA (Avanti, Catalog # 790404), 1% PEG5000-PE (Avanti, Catalog # 880230, and 0.5% atto390-DOPE (ATTO-TEC GmbH, Catalog # AD 390–161. Bilayers were built and proteins coupled as described above. The concentration of CD19 was scaled appropriately to account for the increased surface area of the larger beads.

### Bead engulfment assay

12 to 16 hr prior to imaging, 2.5×10^4^ J774A.1 macrophages expressing the appropriate CAR-P or control construct were plated in a 96-well glass bottom MatriPlate (Brooks, Catalog # MGB096–1–2-LG-L). To assess engulfment, 0.5 × 10^6^ CD19 or CD22-ligated beads were added to each well. Engulfment was allowed to proceed for 45 min at 37ΰC incubator with CO2. Cells were then imaged as described below.

### Bites assay - J774A.1 macrophages, dendritic cells and NIH 3T3 fibroblasts

On the day of imaging 0.5 × 10^6^ NIH 3T3 fibroblasts, dendritic cells or macrophages and 1.5 million Raji B cells were combined in a 1.5 ml eppendorf tube and pelleted by centrifugation (800 rpm / 68 x g) for 5 min at room temperature. Culture media was decanted to ~100 μl volume and cells were gently resuspended, and allowed to interact in the small volume for 60 min in a 37ΰC incubator with CO2. After incubation cells and beads were diluted to a final volume of 1000 μl and 300 μl of this co-culture plated for imaging in a 96-well glass bottom MatriPlate (Brooks, Catalog # MGB096–1–2-LG-L), and imaged as described below.

### Eating assay read by FACS – J774A.1 macrophages and Raji B cells

20.000 J774A.1 macrophages were plated into 96-well glass bottom MatriPlate (Brooks, Catalog # MGB096–1–2-LG-L) in a final volume of 300 μl complete DMEM (Gibco, Catalog #11965–092) supplemented with 1 x Pen-Strep-Glutamine (Corning, Catalog #30–009-Cl) and 10% fetal bovine serum (FBS) (Atlanta Biologicals, Catalog #S11150H). 52 hrs prior to reading the assay macrophages were stimulated with 500 ng/ml LPS (Sigma, Catalog # L4516). 44 hrs prior to imaging LPS was removed by three sequential gentle washes. After LPS removal 10,000 Rajis expressing Caax-mCherry were added to the well containing stimulated macrophages. The co-culture was incubated for 44 hrs in a 37ΰC tissue culture incubator with 5% CO_2_. After 44 hrs, the remaining number of Raji B cells remaining was analyzed by FACS as follows: 10.000 counting beads were added to the well immediately prior to reading and the cell-counting bead mixture was harvested by pipetting up and down 8x with a p200 pipet. The assay was read on an LSRII (BD Biosciences) and Rajis were identified by the presence of mCherry fluorescence.

### Dendritic cell transduction and differentiation

To produce CAR-P expressing dendritic cells, bone marrow-derived hematopoietic stem cells were lentivirally infected immediately after harvest by spinning with 1:4 concentrated lentivirus:GMCSF-containing media (IMDM supplemented with 10% FBS and PSG) on retronectin (Clontech, Catalog # T100A)-coated plates at 2,200 x g for 45 min at 37°C. Dendritic cells were produced as previously described (Mayordomo et al. 1995) by culturing bone marrow cells for 8–11 days with GMCSF. IL-4 was added 2–3 days before use. Efficient differentiation into CD11c+ dendritic cells was verified by FACS, revealing ≥95% APC-CD11c+ cells (Biolegend, Catalog #N418).

### Antigen cross-presentation assay

The ability of CAR-P to stimulate OTI T cell proliferation was tested using the co-culture assay shown as a schematic in Figure S4A and described previously (Roberts et al. 2016). 10,000 CAR-P transduced CD11c+ dendritic cells transduced and differentiated as above were plated in U bottom 96 well dishes (Falcon, Catalog #353077) and stimulated with 1ug/ml LPS. 12 hrs after LPS stimulation, 40,000 Raji B cells expressing soluble cytosolic ovalbumin (Raji B-OVA) were added to the culture. 24 hrs after Raji B-OVA cell addition, 50,000 OTI CD8+ T cells isolated from lymph nodes of OTI TCR transgenic mice using a CD8+ T cell purification kit (Stemcell, Catalog #19853) and labeled with e670 proliferation dye (Thermo, Catalog #65–0840–85) were added. 72hrs after OTI addition the percent of OTI cells divided was measured by eFluor670 signal using flow cytometry.

### Confocal imaging

All imaging in this study was performed using a spinning disk confocal microscope with environmental control (Nikon Ti-Eclipse inverted microscope with a Yokogawa spinning disk unit). For bead internalization assays, images were acquired using a 40×0.95 N/A air objective and unbiased live image acquisition was performed using the High Content Screening (HCS) Site Generator plugin in μManager^3^. Other images were acquired using either a 100×1.49 N/A oil immersion objective. All images were acquired using an Andor iXon EM-CCD camera. The open source μManager software package was used to control the microscope and acquire all images^3^.

Quantification of whole cell internalization 20,000 J774A.1 macrophages were plated into 96-well glass bottom MatriPlate (Brooks, Catalog # MGB096–1 −2-LG-L). Four hours prior to imaging, the macrophages were stimulated with 500 ng/ml LPS (Sigma, Catalog # L4516). Immediately prior to imaging the LPS-containing media was replaced with Fluobrite DMEM (ThermoFisher Scientific, Catalog # A1896701) containing 10% FBS. 40,000 Raji cells were added to the macrophages and the co-culture was imaged at 5 minute intervals for 12 hours. Because cells moved in and out of the field of view, we selected the cells present after 8 hours of imaging and quantified their B cell eating if they could be followed for four hours or more. Time-lapse analysis was essential to ensure that the B cell appeared viable prior to engulfment by the macrophage. Engulfment of B cells with an apoptotic morphology was not counted as a whole cell eating event.

### Quantification of bites internalization

During live cell image acquisition GFP-positive J774A.1 macrophages or NIH 3T3 cells were selected by the presence of GFP signal. A full z-stack comprising the entire cell was captured using 1 μm steps. All z sections were then manually inspected for internalized Raji B cell material. Cells containing one or more bites of fully internalized Raji B cell material >1 μm in diameter were scored as positive.

### Liposome FRET assay

Experiments were carried out as previously described^2^. Briefly, proteins were purified using a bacterial expression system. All protein components (1 mg/ml BSA, 100 nM tSH2-Syk SNAP-505, 0 to 500 nM His_10_-CD3ζ or His_10_-FcRγ intracellular chain, and 7.2 nM His_10_-Lck Y505F) were mixed into kinase buffer (50 mM HEPES-NaOH pH 6.8, 150 mM NaCl, 10 mM MgCl2, and 1 mM TCEP). Liposomes prepared at the following molar ratios: 74.5 % POPC (Avanti, Catalog # 850457C), 10% DOGS-NTA (Nickel) (Avanti, Catalog # 790404C, 0.5% Rhodamine PE (Avanti, Catalog # 810150C), and 15% DOPS (Avanti, Catalog # 840035C) were added and the mixture was incubated for 40–60 min at room temperature, during which the SNAP-505 fluorescence was monitored at 8 s intervals with 504-nm excitation and 540-nm emission. 1 mM ATP was then injected to trigger Lck mediated phosphorylation of CD3ζ or FcRγ. Injection was followed by 5 s of automatic shaking of the plate, and the fluorescence was further monitored at 8 s intervals for at least 1 hr. Data were normalized by setting the average fluorescence value of the last 10 data points before ATP addition as 100% and background fluorescence as 0%. The final extent of fluorescence quenching (% fluorescence change) at each concentration of receptor was determined using the average of the last 20 data points after ensuring fully equilibrated binding. Nine reactions containing increasing concentrations of CD3ζ and nine reactions containing increasing concentrations of FcRγ were run in parallel. The final % fluorescence change was plotted against FcRγ or CD3ζ concentration. The apparent dissociation constants (Kd) of tSH2-Syk to FcRγ and CD3ζ were calculated by fitting the data with Graphpad Prism 6.0, using the “one site specific binding” model.

### Protein expression, purification, and labeling

The intracellular portion of the FcR γ-chain (aa 45–85, Human FcRγ, Uniprot FCERG_HUMAN) was cloned into a modified pET28a vector containing a His_10_ upstream to the multiple cloning site using BamHI and EcoRI. The intracellular portion of CD3ζ (aa 52–164, Human CD3ζ, Uniprot CD3Z_HUMAN) was also cloned into the His_10_ modified pET28a vector. A Lys-Cys-Lys-Lys sequence, originally present for fluorescent labeling, is also present between His_10_ and CD3ζ in this construct. SNAP-tSH2Syk (aa 1–262) was cloned into a pGEX6 vector using BamHI and EcoRI. His_10_-CD3ζ, His_10_-FcR γ-chain, and GST-SNAP-tSH2Syk were bacterially expressed in BL21 (DE3) RIPL strain of *Escherichia coli* as described previously^2^. His_10_-Lck Y505F was expressed in SF9 cells using the Bac-to-Bac baculovirus system as described previously^2^. All cells were lysed in an Avestin Emulsiflex system. His_10_ proteins were purified by using Ni-NTA agarose (Qiagen, Catalog # 30230) and GST-SNAP-tSH2Syk was purified by using glutathione-Sepharose beads (GE Healthcare, Catalog # 17075601) as described previously^2^. Soluble SNAP-tSH2 Syk was generated by cleaving the GST moiety via the PreScission Protease at 4°C overnight. All proteins were subjected to gel-filtration chromatography using a Superdex 200 10/300 GL column (GE Healthcare, Catalog # 17517501) in HEPES-buffered saline (HBS) containing 50 mM HEPES-NaOH (pH 6.8 for His_10_-CD3ζ, His_10_-FcR γ-chain, and GST-SNAP-tSH2Syk and pH 7.4 for His_10_-Lck Y505F), 150 mM NaCl, 5% glycerol, and 1 mM TCEP. The monomer fractions were pooled, frozen in liquid nitrogen and stored at −80 °C. All gel-filtered proteins were quantified by SDS-PAGE and Coomassie staining, using BSA as a standard. To prepare fluorescently labeled tSH2 Syk, 10 uM SNAP- tSH2 Syk was incubated at a 1:2 ratio with SNAP-Cell 505-Star (NEB, Catalog # S9103S) overnight at 4°C and run over a PD MiniTrap G-25 (GE Healthcare, Catalog # 28–9225–29 AB) column to eliminate excess dye.

### Phosphotyrosine Antibody Staining

To fix and stain preparations described above in bead and bites assays for quantifying enrichment of phosphotyrosine staining, half the media (~150 μl) was gently removed from the imaging well and replaced with 150 μl 6.4% paraformaldehyde solution (prepared from 32% stock, Electron Microscopy Sciences, Catalog # 50980495) in tissue culture grade PBS, pH7.2 (Gibco, Catalog # 20012050). Cells were fixed for 15 minutes in a 37ΰC incubator with CO_2_. After fixation cells were washed 2x with PBS and permeabilized/blocked for 60 min at room temperature in freshly prepared, filter sterilized PBS + 5% FBS + 0.1% w/v saponin (PFS solution). After permeabilization, cells were washed 2x 3 min with PFS solution. Following block, cells were incubated with 1:100 dilution of mouse anti-phosphotyrosine (pTyr) antibody to stain pan-pTyr (Santa Cruz, Catalog # PY20) diluted in PFS solution in the dark for 60 min at room temperature then washed 3x 5 min in PFS solution. Washed cells were incubated with a 1:500 dilution of goat anti-mouse Alexa Fluor 647 antibody (Thermo/Molecular Probes, Catalog # A21236) in PFS solution in the dark for 60 min at room temperature. Wells were then washed 3x 5 min in PFS solution. Cells were covered in 200 μl PBS. If not imaged immediately samples were wrapped in parafilm and foil and stored at 4ΰC prior to microscopy. Phosphotyrosine enrichment at the synapse was calculated by dividing the mean Alexa Fluor 647 signal of a 5 pixel linescan at the synapse with bead or cell by a 5 pixel linescan on the cortex. Each data point represents a single cell, and the graphs reflect pooled results from three biological replicates.

### Ovalbumin Antibody Staining

To fix and stain preparations described above for ovalbumin staining, half the media (~150 pl) was gently removed from the imaging well and replaced with 150 μl 8% paraformaldehyde solution (prepared from 32% stock, Electron Microscopy Sciences, Catalog # 50980495) in tissue culture grade PBS, pH7.2 (Gibco, Catalog # 20012050). Cells were fixed for 10 minutes in a 37ΰC incubator with CO_2_. After fixation cells were washed 2x with PBS and permeabilized/blocked for 60 min at room temperature in freshly prepared, filter sterilized PBS + 0.1% w/v casein + 0.1% w/v saponin (PCS solution). After permeabilization, cells were washed 1×3 min with PCS solution and blocked for 1hr at room temperature in PCS. Following block, cells were incubated with 1:100 dilution of rabbit anti-ovalbumin (OVA) antibody to stain OVA (Thermo/Pierce, Catalog # PA1–196) diluted in PCS solution overnight at 4ΰC. Washed cells were incubated with a 1:200 dilution of goat anti-rabbit Alexa Fluor 647 antibody (Thermo/Molecular Probes, Catalog # A21235) and 3.3 nM 488 phallodin (dissolved at 6.6 μM in methanol) in PCS solution in the dark for 60 min at room temperature. Wells were then washed 3×5 min in PCS solution. Cells were covered in 200 μl PBS and immediately imaged. Ovalbumin signal was quantified as the corrected total cell fluorescence (CTCF). CTCF = Integrated Density – Area of Selected Cell * Mean Fluorescence of 3 Background Readings. Each data point represents a single cell, and the graphs reflect pooled results from three biological replicates.

### Image processing and analysis

All image quantification was done on raw, unedited images. All images in figures were first analyzed in ImageJ, where a single Z-slice at the center of the cell was extracted. The image intensities were scaled to enhance contrast and cropped in Photoshop. For movies, background was subtracted in Fiji using a rolling ball radius of 50 μm and bleach corrected using the Histogram Matching plug in.

### Statistics

All statistical analysis was performed in Prism 6.0 (GraphPad, Inc.). The statistical test used is indicated in each figure legend. Error bars throughout the paper denote 95% confidence intervals of the mean. *** indicates p<0.0001; ** indicates p<0.001 and * indicates p<0.01.

---

**Figure S1:**
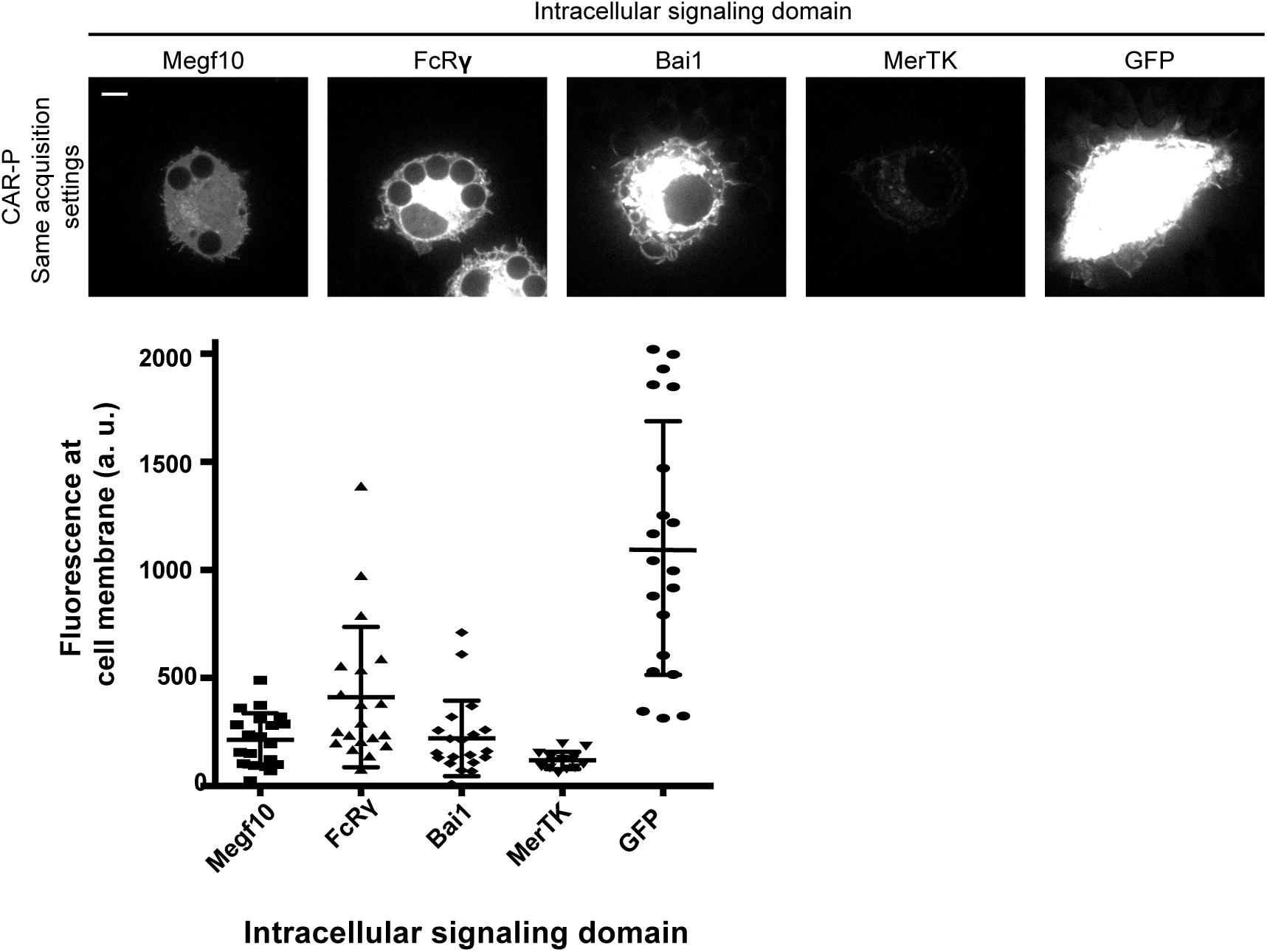
Expression of CAR-P constructs in macrophages. Images of macrophages infected with various αCD19 CAR-P^GFP^ constructs were acquired with identical acquisition settings and scaling to depict differences in expression levels. Fluorescent intensity at the cell cortex of 20 representative αCD19 CAR-P^GFP^-infected macrophages was quantified using the mean intensity of a 2 pixel width linescan at the cell membrane, minus the mean intensity of a linescan immediately adjacent to the cell. The images are the same cells included in Figure 1B and fluorescent intensity was measured from the same macrophages assayed in Figure 1C. The scale bar indicates 5 *μ*m.

**Figure S2:**
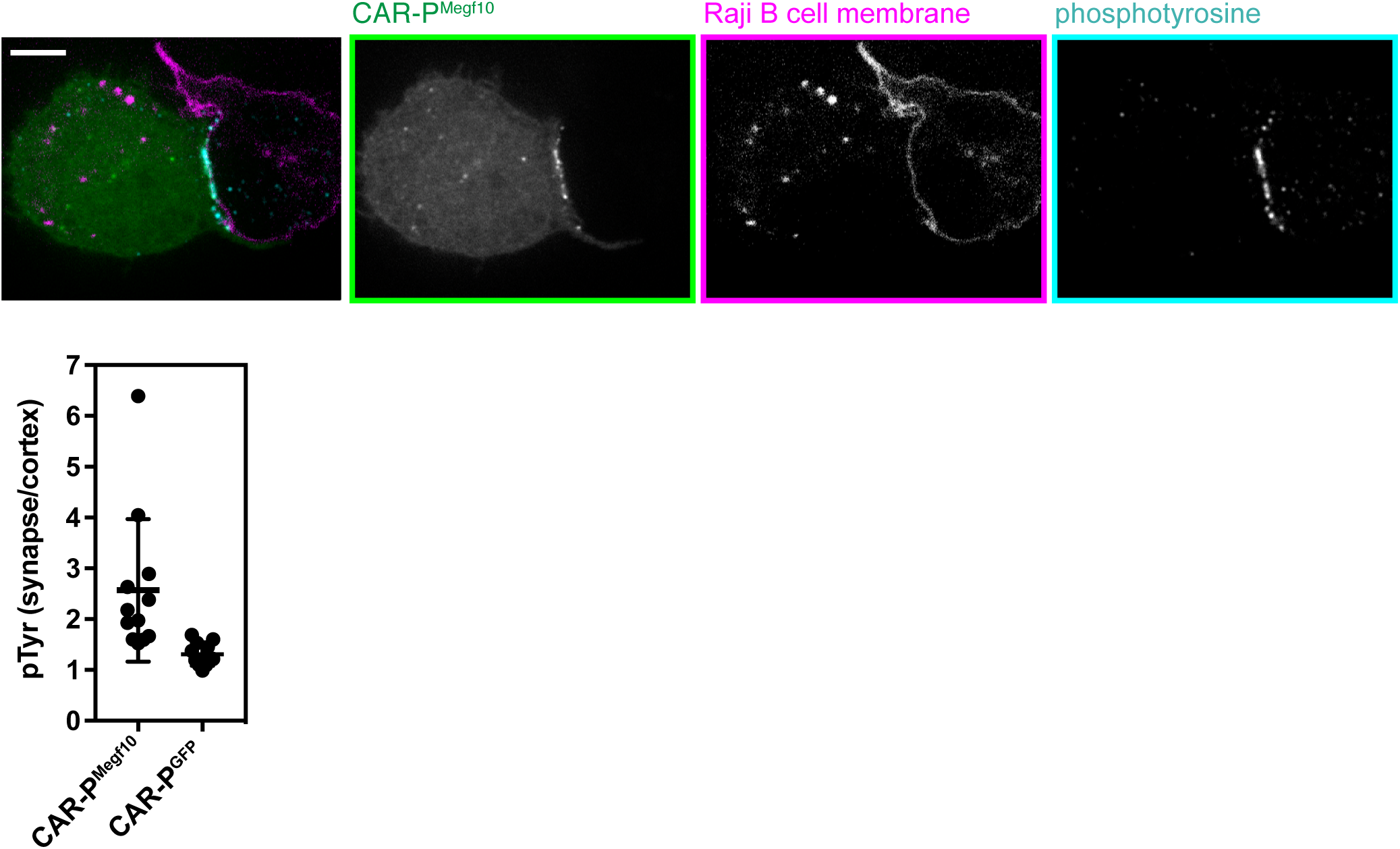
CAR-P localizes with pTyr at synapse with Raji B cell. Phosphotyrosine staining (teal) of macrophages expressing CAR-P^Megf10^ (green) in contact with Raji B cells (cell membrane visualized with mCherry-CAAX, magenta). Below, the enrichment at the synapse is quantified as the mean intensity of a 5 pixel width linescan at the synapse divided by the mean intensity at the adjacent cell cortex for at least 11 sites of contact. Each dot represents one cell-cell synapse, lines represent the mean ± one standard deviation, and the graph is the pooled results o three biological repli-cates. The scale bar indicates 5 *μ*m.

**Figure S3:**
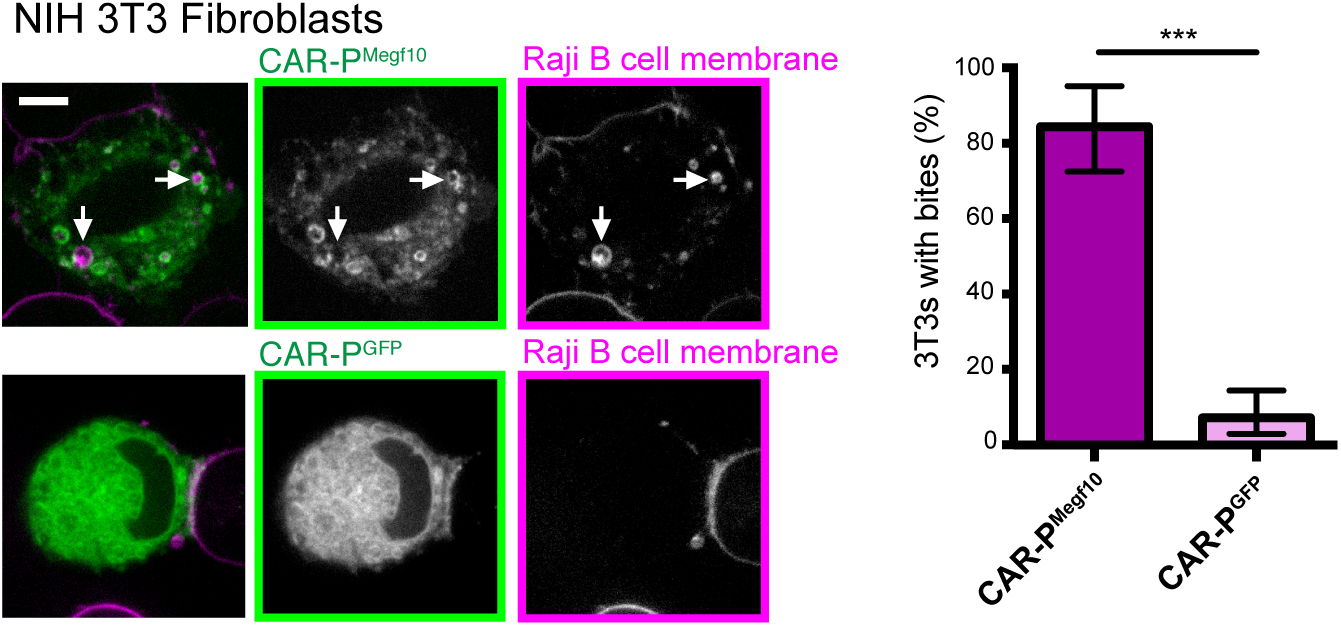
NIH 3T3 cells internalize Raji B cell bites. NIH 3T3 cells expressing the αCD19 CAR-P^Megf10^ (green in merge, left; greyscale, center) engulf pieces of CD19+ Raji B cells (labeled with mCherry-CAAX; magenta in merge, left; greyscale, right). The control αCD19 CAR-P^GFP^-infected 3T3s are shown below. Arrows point to pieces of ingested Raji B cell. The proportion of cells taking at least one bite after 90 min co-incubation is graphed on the left (graphs show the pooled data of three separate experiments; n=111 CAR-P^Megf10^ 3T3 cells and 121 CAR-P^GFP^ 3T3; *** indicates p<0.0001 by two-tailed Fisher Exact Test; error bars denote 95% confidence intervals). Bites are defined as a fully internalized piece of mCherry-labeled material >1 *μ*m in diameter.

**Figure S4:**
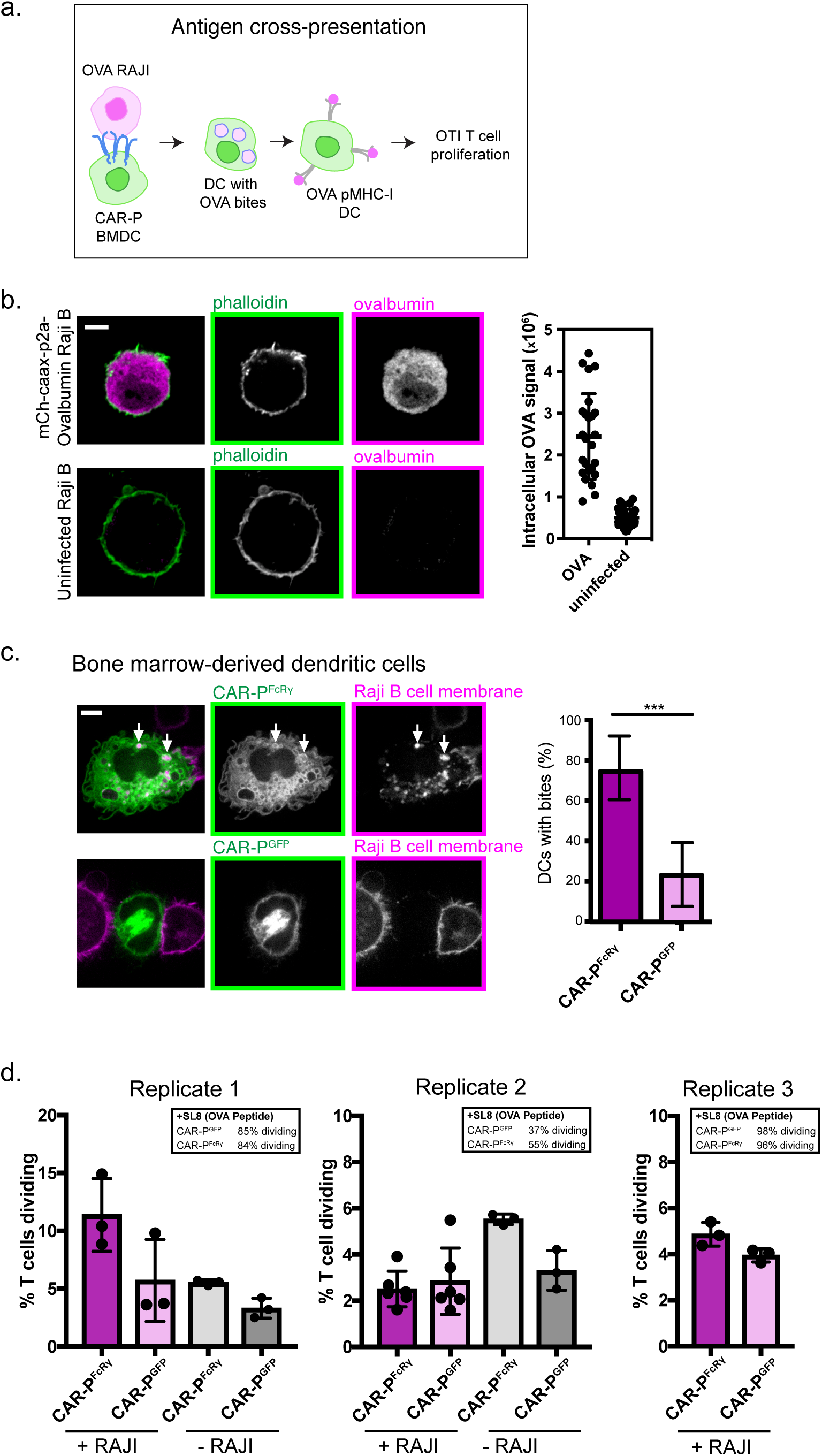
CAR-Ps promote internalization of cancer antigen(A) Schematic of antigen internalization and cross-presentation assay. CAR-P expressing Bone Marrow Derived Dendritic Cells (BMDC) were differentiated using GM-CS. CAR-P BMDC were incubated with Raji B cells expressing soluble ovalbumin (OVA). DC with OVA bites (internalized antigen) were then incubated with OTI T cells (OVA specific CD8+ T cells) and OTI proliferation assessed as a measure of T cell stimulation. Results from each step of this assay are shown in sequence in (B), (C), and (D). (B) Ovalbumin staining (magenta) in Raji B cells infected with Caax-mCherry-p2a-Ovalbumin lentivirus (OVA) and uninfected controls (uninfected) shows robust OVA expression in infected cells. At right the intracellular OVA signal is plotted as corrected total cell fluorescence (CTCF) for the ovalbumin channel. Each dot represents the CTCF of one cell; n=26 cells OVA, n=33 cells (uninfected); lines represent the mean ± one standard deviation, and the graph is the pooled results of three biological replicates. The scale bar indicates 5 μm. (C) Bone marrow-derived dendritic cells expressing the CAR-P^FcRγ^ (top panel, green in merge, left; greyscale, center) engulf pieces of CD19+ Raji B cells (labeled with mCherry-CAAX; magenta in merge, left; greyscale, right). The control αCD19 CAR-P^GFP^-infected dendritic cells are shown below. Arrows point to pieces of ingested Raji B cell. The proportion of cells taking at least one bite after 90 min co-incubation is graphed on the right of images. Graphs show the pooled data of two separate experiments; n=28 CAR-P^FcRγ^ dendritic cells and n=33 CAR-P^GFP^ dendritic cells; *** indicates p<0.000l by two-tailed Fisher Exact Test; error bars denote 95% confidence intervals. Bites are defined as a fully internalized piece of mCherry-labeled material >1 μm in diameter. (D) OTI T cell proliferation after 72hr incubation with CAR-P transduced CD11c+ dendritic cells. +/- RAJI below the x-axis indicates whether Raji-OVA B cells were added to CAR-P transduced dendritic cells prior to OTI addition. To measure proliferation, T cells were uniformly stained with eFluor670 dye on day 0, and proliferation was measured by dilution of the cell-bound dye. Graphs show the mean ± SD of three independent biological replicates. Data points are values for individual wells of differentiated CD11c+ dendritic cells. Boxed data indicate the mean % T cells dividing when dendritic cells were pulsed with SL8 (OVA) peptide, which directly binds to MHC without undergoing cross presentation. If dendritic cell differentiation was successful, the pulsed dendritic cells should be capable of inducing robust OTI proliferation. Sample sizes were selected to match previous studies that were able to detect robust T cell stimulation (Roberts et al. 2016).

Video 1: CAR-P^Megf10^ macrophage engulfs silica beads

A macrophage infected with αCD19 CAR-P^Megf10^ (green) engulfs 5 μm silica beads coated in a supported lipid bilayer (labeled with atto647, magenta) and ligated to his-tagged CD19 extracellular domain. The field of view is 43 × 43 μm. The movie is a maximum intensity projection of 17 z-planes acquired at 0.5 μm intervals. Z-stacks were acquired every 30 sec for 30 min and time is indicated in the bottom right.

Video 2: CAR-P^Megf10^ macrophage engulfs bites of a Raji B cell

A macrophage infected with αCD19 CAR-P^Megf10^ (green) engages with a Raji B cell (labeled with mCherry-CAAX). The field of view is 43 × 43 μm. The movie is a maximum intensity projection of 7 z-planes acquired at 1 μm intervals. Images were acquired every 20 sec for 30 min and time is indicated in the bottom right.

Video 3: CAR-P^FcR^* macrophage engulfs a Raji B cell.

A macrophage infected with αCD19 CAR-P^FcRγ^ (green) engages with a Raji B cell (labeled with mCherry-CAAX). The field of view is 53×53 μm. Images were acquired every 5 min. Time is indicated in the bottom right.

Video 4: CAR-P^tandem^ macrophage engulfs a Raji B cell.

A macrophage infected with αCD19 CAR-P^tandem^ (green) engages with a Raji B cell (labeled with mCherry-CAAX). The field of view is 53×53 μm. Images were acquired every 5 min. Time is indicated in the bottom right.

